# Proteomic identification of Axc, a novel beta-lactamase with carbapenemase activity in a meropenem-resistant clinical isolate of *Achromobacter xylosoxidans*

**DOI:** 10.1101/191403

**Authors:** Frank Fleurbaaij, Alex A. Henneman, Jeroen Corver, Cornelis W. Knetsch, Wiep Klaas Smits, Sjoerd T. Nauta, Martin Giera, Irina Dragan, Nitin Kumar, Trevor D. Lawley, Aswin Verhoeven, Hans C. van Leeuwen, Ed J. Kuijper, Paul J. Hensbergen

**Author notes:** Correspondence to: P.J. Hensbergen, Center for Proteomics and Metabolomics, Leiden University Medical Center, PO Box 9600, 2300 RC Leiden, The Netherlands.

## Abstract

The development of antibiotic resistance during treatment is a threat to patients and their environment. Insight in the mechanisms of resistance development is important for appropriate therapy and infection control. Here, we describe how through the application of mass spectrometry-based proteomics, a novel beta-lactamase Axc was identified as an indicator of acquired carbapenem resistance in a clinical isolate of *Achromobacter xylosoxidans*.

Comparative proteomic analysis of consecutively collected susceptible and a resistant isolates from the same patient revealed that high Axc protein levels were only observed in the resistant isolate. Heterologous expression of Axc in *Escherichia coli* significantly increased the resistance towards carbapenems. Importantly, direct Axc mediated hydrolysis of imipenem was demonstrated using pH shift assays and ^1^H-NMR, confirming Axc as a legitimate carbapenemase. Whole genome sequencing revealed that the susceptible and resistant isolates were remarkably similar.

Together these findings provide a molecular context for the fast development of meropenem resistance in *A. xylosoxidans* during treatment and demonstrate the use of mass spectrometric techniques in identifying novel resistance determinants.

## Introduction

Development and spread of antibiotic resistance by pathogenic microorganisms is an increasing healthcare problem. Moreover, certain resistance determinants spread readily ^1,2^, while the introduction of novel antibiotics is lagging behind. Several clinically important classes of antimicrobials such as the beta-lactams, target the bacterial cell wall ^3^. Resistance to beta-lactams can be mediated by beta-lactamases that are capable of hydrolysing the beta-lactam ring. Following the initial introduction of penicillin, second and third generation beta-lactams have been developed which, in turn, triggered the selection of beta-lactamases with broader specificities. Carbapenem treatment is often used as a last resort, since extended-spectrum beta-lactamases (cephalosporinases) are becoming more prevalent in Gram-negative bacteria. The emergence and spread of carbapenemases such as class A KPC ^4^, a number of metallo beta-lactamases ^5,6^ (class B: IMP, VIM, NDM) and class D oxacillinases such as OXA-48 ^7^, in combination with other resistance mechanisms ^8^, can jeopardize carbapenem efficacy, leaving little or no treatment options for patients.

*Achromobacter xylosoxidans* is a rod shaped aerobic non-fermentative Gram-negative bacterium. It is widespread in the environment and generally considered as an opportunistic pathogen. Chronic infections with *A. xylosoxidans* are problematic in cystic fibrosis patients ^9,10^ but reported prevalence numbers vary greatly (3-30%) ^11,12^. Moreover, bacteremia as a result of *A. xylosoxidans* can occur in immunocompromised patients ^13^. *A. xylosoxidans* is notorious for its intrinsic high level of resistance, especially towards penicillins and cephalosporins ^14–16^. In general, carbapenem resistance in *A. xylosoxidans* is not widespread and as a result meropenem treatment is routinely applied, even in the case of recurring infections ^17,18^. Though carbapenem resistance is observed, specifically for meropenem ^19^, there are few reports on the mechanism of carbapenemase resistance in *A. xylosoxidans*. Notable exceptions are the plasmid-encoded carbapenemase IMP ^20,21^ and the chromosomally encoded class D beta-lactamase OXA-114 ^16^, that show low level carbapenemase activity. A comparative genomic exploration of two *A. xylosoxidans* isolates revealed many genes that could be involved in drug resistance, such as efflux pumps and β-lactamases. However, most of these genes were conserved between carbapenem susceptible and resistant strains, highlighting the difficulty in translating genomic data to the observed resistant phenotypes ^22^.

In this study, two clinical isolates of *A. xylosoxidans* from an immunocompromised patient with pneumonia were investigated. The initially cultured isolate from the respiratory tract was susceptible to meropenem and treatment was started accordingly. However, a subsequent meropenem resistant isolate was obtained from a blood culture after treatment failure. Since PCR analysis was negative for known carbapenemases, we performed a proteomic analysis which revealed the novel beta-lactamase Axc as highly abundant in the meropenem-resistant, but not in the susceptible isolate. Axc expression led to an increase of minimal inhibitory concentrations for carbapenems when introduced in a susceptible *Escherichia coli* strain and direct carbapenemase activity of Axc was demonstrated using *in vitro* imipenem conversion assays. Interestingly, the resistant as well as the susceptible clinical isolates are genetically almost identical, emphasizing the importance of mass spectrometry as a technique to investigate carbapenem resistance in *A. xylosoxidans*.

## Results

### Development of meropenem resistance in *Achromobacter xylosoxidans* during treatment

A 65-year old patient, diagnosed with chronic lymphocytic leukemia in 1989, underwent a non-myeloablative stem cell transplantation in July 2014. In August 2014, the patient developed neutropenic fever and pneumonia due to an infection with *A. xylosoxidans*. The initial antibiogram (Supplemental Table 1) revealed a multi-resistant character, as is commonly found for *A. xylosoxidans*, but the isolate was susceptible to meropenem. Hence, meropenem treatment was initiated (4 dd 1 gram intravenously). During treatment the patient developed a pneumothorax and died from septic shock 4 days later. At this point the patient had been treated with meropenem for six days, as well as vancomycin and antifungal therapy with liposomal amphotericin B. An antibiogram of a blood culture from a sample taken one day before the patient’s death revealed a meropenem-resistant *A. xylosoxidans* phenotype (Supplemental Table 1). Subsequently, pure cultures from the first and second clinical isolate were prepared (AchroS and AchroR, respectively). In line with the antibiogram analysis described above, Etests showed that both isolates were resistant to imipenem (MIC > 32 mg/L for both) but differed in their susceptibility towards meropenem (MIC of 0.0094 mg/L (first isolate, AchroS) and 2 mg/L (second isolate, AchroR), respectively). An in-house multiplex PCR assay (based on a published PCR ^34^) failed to detect common carbapenemases (KPC, IMP, VIM, NDM and OXA-48, data not shown), suggesting that the change in resistance is not mediated by these enzymes. This finding prompted us to perform a comparative proteomic analysis of the two isolates to attempt to identify the meropenem resistance mechanism.

**Table 1:** Effect of Axc expression in *E. coli* on the susceptibility towards imipenem and meropenem. Minimal inhibitory concentrations (MICs) of imipenem and meropenem for *E.coli* CA43 expressing Axc (*E. coli*_ Axc), or PPEP-1 (*E. coli_*control) and *Klesbsiella pneumoniae* expressing KPC carbapenemase (KPC). N.D: Not detectable. IPTG: isopropyl β-D-1-thiogalactopyranoside.

| Strain | MIC (mg/L) |  |  |  |
| --- | --- | --- | --- | --- |
|  | Imipenem |  | Meropenem |  |
|  | - IPTG | + IPTG | -IPTG | + IPTG |
| *E. coli*_Axc<sup>a</sup> | 0,5 | 4 | 1,563 | 12,5 |
| *E. coli*_control<sup>b</sup> | 0,38 | N.D. | 0,391 | 0,391 |
| KPC<sup>c</sup> | >32 | >32 | >12,5 | >12,5 |

<sup>a</sup>JC107: *Escherichia coli* C43/pET21B-Axc
<sup>b</sup>JC108: *Escherichia coli* C43/pET21B/PPEP-1 (Pro-Pro endopeptidase 1)
<sup>c</sup>JC113: Carbapenemase-positive *Klebsiella pneumoniae*

### Comparative proteomic analysis shows differential levels of the beta-lactamase Axc

Protein extracts of the meropenem-susceptible and resistant *A. xylosoxidans* isolates (AchroS and AchroR) were first analysed by SDS-PAGE. Since no major visual differences were observed (Supplemental Figure 1A), all bands were excised and processed for a bottom-up LC-MS/MS proteomic analysis. In total, 2290 unique proteins were identified, of which 1517 proteins were common to both isolates, while 226 and 537 were only found in the resistant and susceptible isolates, respectively. For a semi-quantitative analysis, the spectra assigned to peptides belonging to a certain protein were counted and compared between the two different isolates (Supplemental Figure 1B). Of the uniquely observed proteins, most are proteins with low spectral counts (often single peptide identifications), likely resulting from sampling bias of low abundant proteins. One protein was observed with 100 spectra in the resistant isolate (AchroR) but none in the susceptible isolate (AchroS). This protein, hereafter called Axc (for *<u>A</u>chromobacter <u>x</u>ylosoxidans* <u>c</u>arbapenemase, GenBank ID: MF767301), is a putative PenP class A beta-lactamase (COG2367/pfam13354).

**Figure 1:**
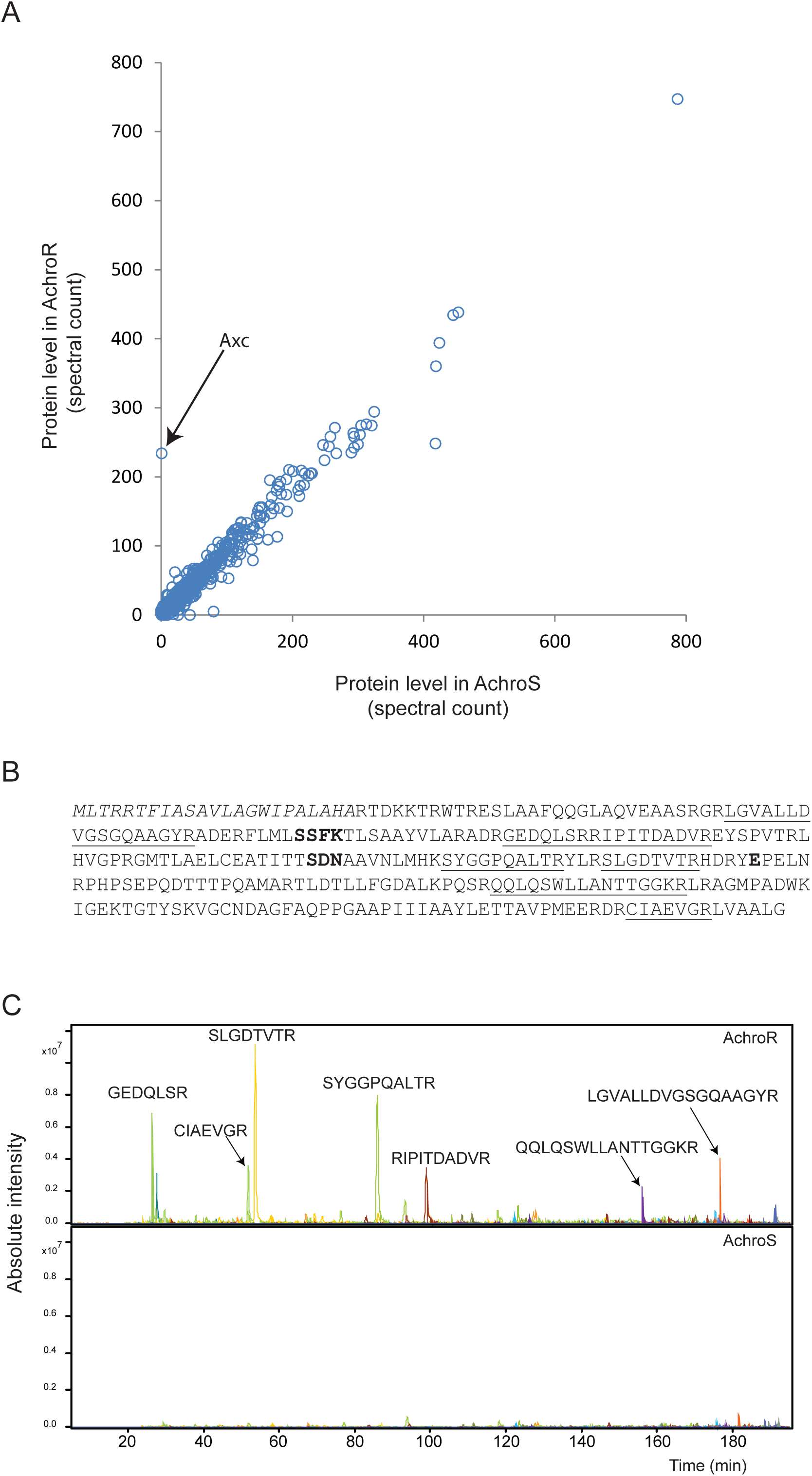
Comparative proteomic analysis of meropenem resistant and susceptible *Achromobacter xylosoxidans* clinical isolates. A: Tryptic digests of protein extracts of the meropenem resistant (AchroR) and susceptible (AchroS) isolate were analysed by LC-MS/MS. Spectra were assigned to peptides based on database searching. Identified spectra were then assigned to the corresponding proteins and the total number of spectra per protein were counted. Each circle represents one protein with the number of spectra observed in the resistant and the susceptible isolate. Hence, proteins on the diagonal were observed in similar counts in both isolates. Axc (arrow), a classA PenP-family beta-lactamase, is the most prominent outlier. B: The full amino acid sequence of Axc, with the peptides identified by LC-MS/MS analysis underlined. Conserved residues from serine beta-lactamases, Ser-X-X-Lys, Ser-Asp-Asn and the active site Glu, are in bold (37). C: Extracted ion chromatograms of *m/z* values corresponding to tryptic peptides of Axc in the meropenem resistant isolate (AchroR, upper panel) and susceptible isolate (AchroS, lower panel). The corresponding tryptic peptides are indicated above the corresponding peaks.

To confirm that Axc is highly abundant in the resistant isolate in comparison to the susceptible isolate, a second proteomic analysis was performed on whole cell protein extracts that were digested in-solution and analysed without any prior fractionation. Spectral count analysis resulted in a cumulative quantification of 1356 different proteins, of which 987 were found in both isolates, but 202 and 167 were uniquely quantified in the resistant and susceptible isolates, respectively. The spectral count plot reflects the high similarity of the two clinical isolates, with the vast majority of the proteins distributed along the diagonal (Figure 1A). In accordance with the data described above, Axc is the most prominent outlier in the resistant clinical isolate. A number of Axc tryptic peptides (Figure 1B) is clearly visible in the LC-MS/MS analysis of the resistant but not the susceptible isolates (Figure 1C). Of note, cells used for these analyses were grown in the absence of meropenem, so the high level of Axc in the resistant isolate is independent of antibiotic pressure.

### *Axc* is present in meropenem resistant and susceptible *A. xylosoxidans* isolates

To investigate whether *A. xylosoxidans* acquired the *axc* gene in the course of the treatment, we performed a PCR for the *axc* open reading frame on both the resistant and susceptible isolates AchroS and AchroR. This analysis demonstrated that *axc* is present in both clinical isolates (Supplemental Figure 2). Moreover, Sanger sequence analysis demonstrated that the sequence of *axc* is identical in both isolates (our unpublished observations), indicating that the observed meropenem resistance is not likely caused by an alteration of protein function.

**Figure 2:**
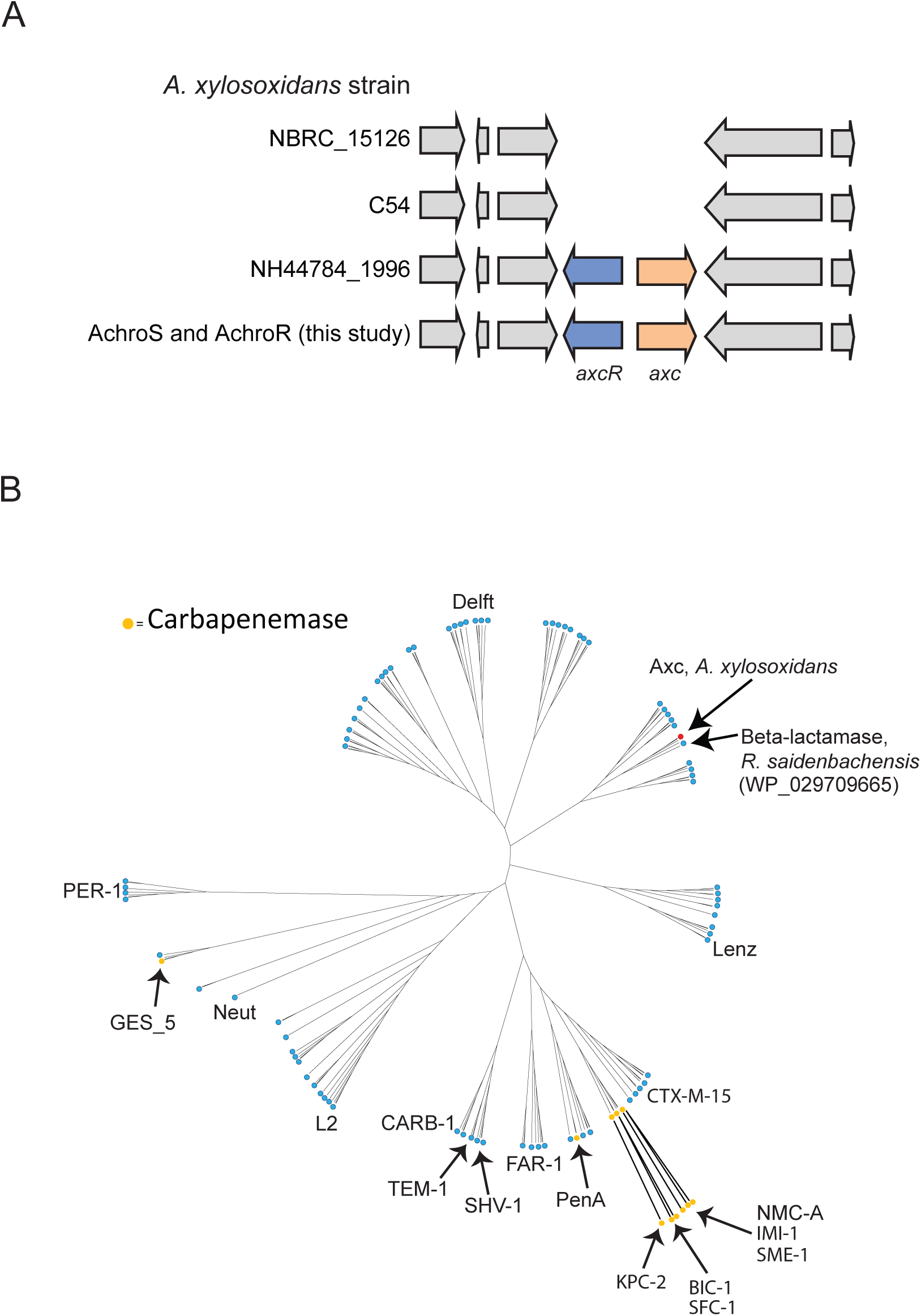
Genomic context of *axc* in *Achromobacter xylosoxidans* strains and comparison of Axc with other class A beta-lactamases. A: *Axc* (a putative PenP class A beta-lactamase, 1.17 e-^54^) and the gene encoding its putative transcriptional repressor (*axcR*), were found in both clinical isolates (AchroS and AchroR). Three other fully sequenced genomes of *Achromobacter xylosoxidans* were examined for the presence of *axc*; NH44784_1996 (Genbank identifier NC_021285.1), C54 (Refseq assembly GCF_000186185.1) and NBRC_15126/ATCC27061 (Genbank chromosome CP006958.1). Only within the strain NH44784_1996 (used as a reference to search our proteomics data), *axc* and the putative regulator are also present. B: Unrooted cladogram obtained for 176 class A beta-lactamases including Axc. The class A β- lactamase protein sequences of Gram negative bacteria were obtained by querying the refseq_protein database using Blastp (<1e-10, http://blast.ncbi.nlm.nih.gov/blast/Blast.cgi) and a consensus β-lactamase-alignment (37). Duplicate sequences and sequences causing a strong overrepresentation of branches produced in the tree were removed. Names of known carbapenames (orange) and names of identifier of branches (37) are indicated. Axc is indicated with a red dot. The closest homologue to Axc, a β-lactamase from *R. saidenbachensis* (WP_029709665), is also indicated.

A database search revealed that the *axc* gene is not present in all *A. xylosoxidans* strains (Figure 2A). Like in our clinical isolates, *axc* is present in the NH44784_1996 strain (Genbank identifier NC_021285.1) ^35^ but not in the strains NBCR 15126/ATCC27061 (Genbank chromosome CP006958.1) and C54 (Refseq assembly GCF_000186185.1) for instance. In those strains that contain *axc*, the gene is located next to a putative LysR-type transcriptional regulator (Pfam 03466), hereafter *axcR* (for *<u>axc</u>-*associated <u>r</u>egulator). Additional PCR and Sanger sequencing experiments verified that the intergenic regions in AchroS and AchroR are identical, However, they differ at two positions with the intergenic region in strain NH44784_1996 (Supplemental Figure 2B).

Our results show a high degree of similarity between the meropenem resistant and susceptible strain, raising the possibility that these two strains represent a clonal complex. To further explore the relatedness of both isolates, whole genome sequencing (WGS) analysis was performed. This showed that both patient isolates are highly similar, with only one single-nucleotide polymorphism (SNP) within the gene encoding AxyZ ^36^. This SNP was confirmed by PCR and subsequent Sanger sequencing, ruling out the possibility that this was an artefact of the assembly procedure. Moreover, *axc* and its putative regulator *axcR* were found to be located in the same genomic region as in NH44784_1996 strain (Figure 2A). Overall, the genome sequences suggest that both isolates are clonal, and that the meropenem resistance evolved within the same strain during the course of treatment.

To demonstrate how Axc is related to other class A beta-lactamases, we compared the Axc amino acid sequence with the sequence of another 176 representatives of this family using an alignment consensus based on a report by Walther-Rasmussen and Hoiby ^37^. The resulting unrooted cladogram shows that Axc is most closely related to a class A beta-lactamase of *Rhodoferax saidenbachensis* (WP_029709665, Figure 2B). Only a limited number of class A beta-lactamases have activity towards carbapenems, but none of these cluster with the Axc sequence (Figure 2B). Our data therefore suggest a novel function for the PenP family of beta-lactamases (COG2367) to which Axc belongs.

### Functional characterisation of Axc

To establish whether Axc indeed has activity towards carbapenems, Axc was expressed in a heterologous host and hydrolysis of carbapenems was measured indirectly and directly.

The *E. coli* C43 strain, suitable for the expression of toxic proteins ^30^, is susceptible to carbapenems. We generated a derivative of C43 that allows for IPTG-dependent expression a plasmid-based copy of *axc* (*E. coli*_Axc (JC107)). Susceptibility testing for imipenem and meropenem showed that the MICs for these carbapenems increased 8-fold, following induction of *Axc* expression (Table 1). Though these levels were lower than for the positive control (KPC) for our assay, they were specific for Axc as the expression of an unrelated protein (PPEP-1 ^31^) did not lead to an increase in MIC values (Table 1). Thus, expression of Axc is correlated to resistance towards carbapenems.

To directly demonstrate carbapenemase activity, hydrolysis of imipenem was monitored *in vitro* through colorimetric assays (Figure 3A) and ^1^H-NMR (Figure 3B). Imipenem hydrolysis results in the formation of a carboxylic acid, and monitoring the accompanying pH drop colorimetrically is a well-established method for the detection of carbapenemase activity ^32,38^. Indeed, hydrolysis of imipenem was readily observed using KPC cells (without IPTG). Consistent with our previous results, imipenem hydrolysis was observed for *E. coli* cells harbouring the Axc expression plasmid (*E. coli*_Axc) grown in the presence, but not in the absence, of IPTG. As before, these results were specific for Axc, as induction of PPEP-1 expression (*E.coli*_control) did not result in imipenem hydrolysis (Figure 3A). In a parallel assay, ^1^H-NMR was used to directly observe the opening (hydrolysis) of the lactam ring in imipenem (Figure 3B). The chemical shifts of the peaks change as a result of this hydrolysis, with the protons closest to the ring opening undergoing the largest change. The H-6 proton of imipenem generates an adequately resolved multiplet at 3.42 ppm that decreases in intensity upon hydrolysis. Concomitantly, the doublet generated by the protons of the methyl group (H-9) move upfield, resulting in a decrease of the doublet at 1.3 ppm. After 10 minutes incubation of bacterial cells with imipenem, hydrolysis of imipenem was observed with *E. coli*_Axc grown in the presence, but not in the absence, of IPTG. Hydrolysis was also apparent for KPC, but not for *E. coli* cells expressing PPEP-1 (*E.coli*_control)), even after long incubation times (10h). Under these conditions, imipenem hydrolysis was also observed for samples with *E. coli* cells harbouring the Axc expression plasmid grown in absence of IPTG, due to leaky expression from the inducible promoter.

**Figure 3:**
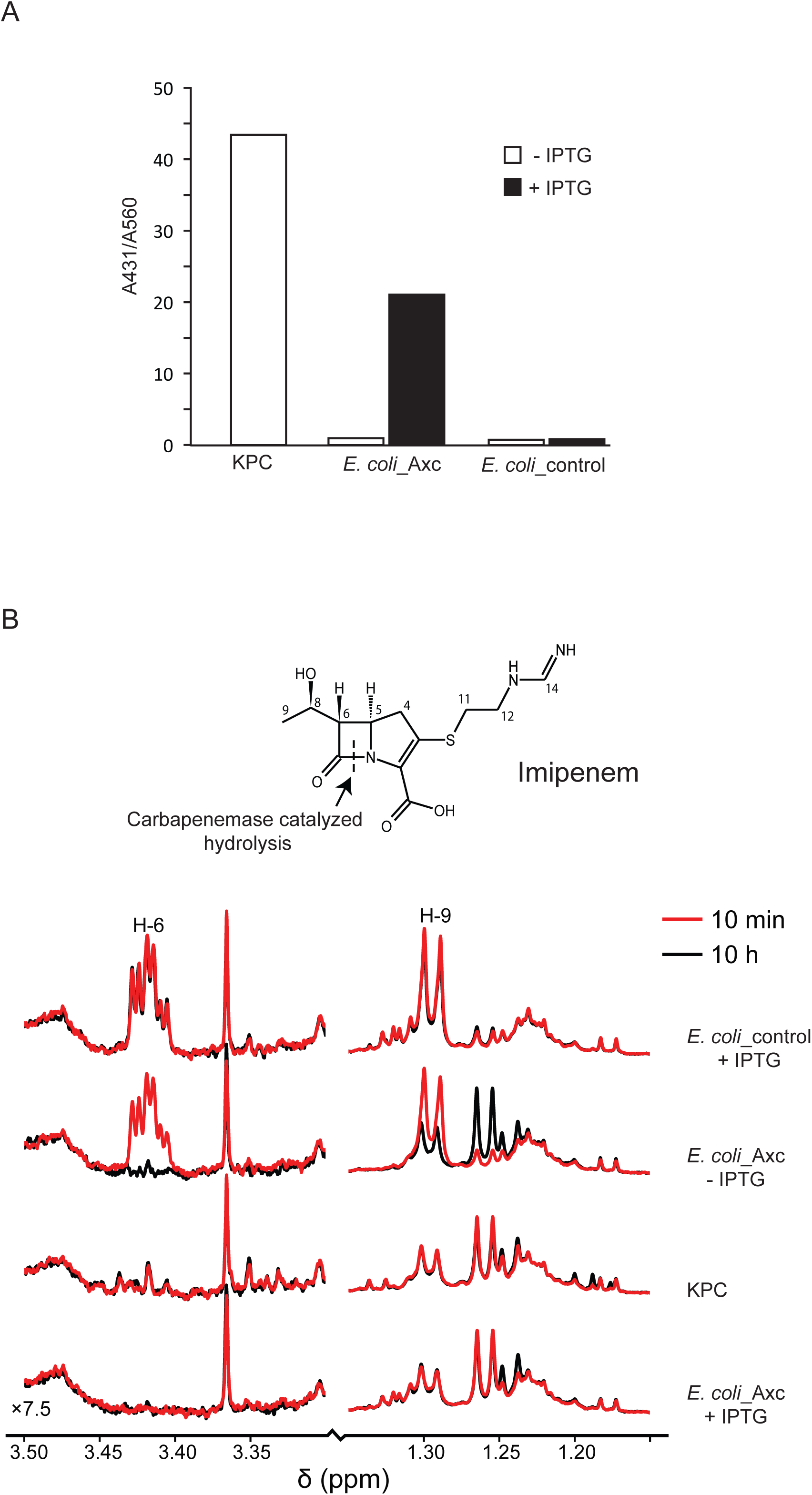
Axc has carbapenemase activity. A: Axc was expressed in *E. coli* and cell extracts were tested for the ability to hydrolyse imipenem. Bars show the A431/A560 ratio, which is a measure for the shift in pH due to imipenem hydrolysis. B: NMR-based identification of imipenem hydrolysis by Axc. The structure of imipenem is shown, with the proton numbering used in the spectrum. The line color indicates the incubation time (red, 10 min at room temperature or black, 10 hrs at 6 °C). Imipenem hydrolysis is accompanied by the loss of the H-6 multiplet at 3.4 ppm, and a shift in the H-9 doublet, resulting in a decrease of the doublet at 1.3 ppm. Strains used (see also Table 2): KPC (JC113): Carbapenem resistant *Klebsiella pneumoniae*. *E.coli* Axc (JC107): *E. coli* strain C43(DE3), containing plasmid pET21-Axc; *E.coli*_control (JC108): *E. coli* strain C43, containing plasmid pET21-PPEP-1 (Pro-Pro endopeptidase 1). IPTG: isopropyl β-D-1-thiogalactopyranoside

Taken together, our data establish that Axc has carbapenemase activity.

**Table 2:**
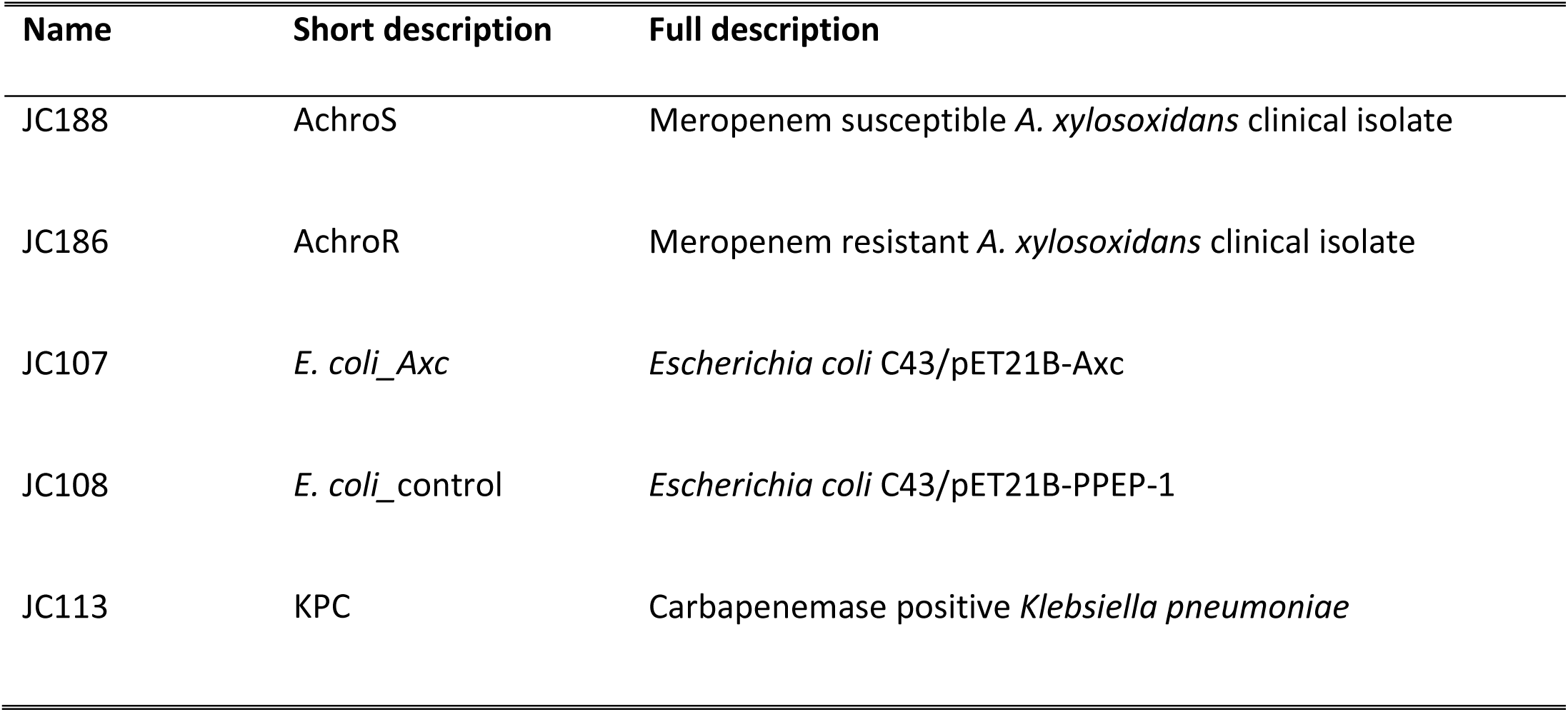
Strains used in this study. Axc:*<u>A</u>chromobacter <u>x</u>ylosoxidans* <u>c</u>arbapenemase. PPEP-1: Pro-Pro endopeptidase 1 (31)

| Name | Short description | Full description |
| --- | --- | --- |
| JC188 | AchroS | Meropenem susceptible *A. xylosoxidans* clinical isolate |
| JC186 | AchroR | Meropenem resistant *A. xylosoxidans* clinical isolate |
| JC107 | *E. coli*_Axc | *Escherichia coli* C43/pET21B-Axc |
| JC108 | *E. coli*_control | *Escherichia coli* C43/pET21B-PPEP-1 |
| JC113 | KPC | Carbapenemase positive *Klebsiella pneumoniae* |

## Discussion

In this paper we identified a new resistance mechanism that explained the difference in meropenem susceptibility of two clinical isolates of *A. xylosoxidans* that were collected within two weeks during treatment. Using a combination of comparative proteomic analyses and functional assays, we have shown that the class A beta-lactamase Axc is highly abundant in the meropenem resistant isolate in comparison to the susceptible isolate and that Axc has carbapenemase activity.

Detection of carbapenemases from sequence data is challenging. First, carbapenemases belong to different subgroups of beta-lactamases which have probably evolved by convergent evolution. Although sequence identities are moderate, most of the class A carbapenemases have a disulphide bridge between Cys-69 and Cys-238, but this is dispensable for activity against carbapenems ^39^ and our data show that Axc does not contain these residues. Thus, the presence of these cysteine residues is no guaranteed predictor for carbapenemase activity. Next, some studies also revealed a mechanism where beta-lactam trapping, without actual degradation, can be involved in resistance towards carbapenems when levels are sufficiently high. In such cases, concomitant loss of porins is often observed ^40,41^. Thus, it is crucial to determine whether a beta-lactamase actually induces hydrolysis of carbapenems. Our NMR and pH-shift analyses data clearly demonstrate Axc-mediated opening of the beta-lactam ring in imipenem. Finally, many unexplained mechanisms of carbapenem resistance remain. For instance, a recent study demonstrated plasmid derived carbapenem resistance in *Klebsiella pneumonia* strains which could not be explained by the most common carbapenemases found (KPC-type). Even though none of the plasmid-encoded genes were obvious candidates for the observed resistance towards carbapenems, several TEM-homologs were detected ^42^. Though the prediction of the activity of a certain beta-lactamase against carbapenems is not straightforward studies such as the present one highlight that mass spectrometry approaches can be used to gain insight in the mechanism of action and role of specific proteins in the observed phenotypes.

We do not know whether the high level of Axc is the only mechanism conferring meropenem resistance to our isolate of *A. xylosoxidans*. When expressed from an inducible promoter, Axc confers moderate resistance to carbapenems to *E. coli*; MIC values compared to the KPC strain suggest that Axc has a lower efficiency than KPC, but may also indicate lower overall levels of expression. Differing efficiencies in carbapenemases are well documented, to the point where the activity of a specific class, such as OXA-48, is difficult to detect but of great clinical importance ^43^. Full biochemical characterization of Axc, including kinetic experiments, is subject to further study. Such experiments, in combination with crystallography analysis, will provide more insight in the activity of Axc against different beta-lactams and could resolve the structural characteristics of the binding pocket which facilitates its activity towards carbapenems. We note, however, that two other changes in the antibiogram between the meropenem susceptible and resistant isolates involve beta-lactam antibiotics. Augmentin (amoxicillin/clavulanate) resistance changed from intermediate (8 mg/L) to resistant (>32 mg/L), and piperacilline/tazobactam from susceptible (<=4 mg/L) to intermediate resistance (8 mg/L). This suggests that Axc may have a broad substrate specificity and is insensitive to inhibition by clavulanate and tazobactam. Strikingly, both the meropenem-susceptible and meropenem-resistant isolates were resistant to imipenem (MIC values higher than the maximum concentration tested (32 mg/L)). This indicates that, notwithstanding the activity of Axc towards imipenem as presented here, imipenem resistance in the clinical isolates is not dependent on Axc. Differences in the sensitivities towards different carbapenems results from the chemical differences between the individual drugs ^44^ and are often linked to the differential permeability of the outer cell membrane ^45,46^.

The regulatory mechanism leading to higher levels of Axc expression are unclear. Sequence analyses showed that the meropenem susceptible and resistant *A. xylosoxidans* clinical isolates are highly similar, with no differences in the *axc* promotor and coding sequence, nor in its putative regulator AxcR and the *axc*-*axcR* intergenic region. The only SNP we identified is located in the gene encoding AxyZ, the TetR-type repressor of the *axyXY-oprZ* operon ^47^. This leads to an amino acid substitution (V29G) in a region of AxyZ that is involved in DNA binding in other members of TetR family ^48^. AxyX, AxyY and OprZ form an efflux pump of the resistance-nodulation division (RND) family and are predominantly found in aminoglycoside resistant *Achromobacter* species ^49–51^. A recent paper showed higher expression levels of *axyY* in *A. xylosoxidans* strains containing AxyZ_Gly29, suggesting that this mutation leads to reduced repression of AxyZ ^52^. Closer inspection of our proteomics data indicates that also AxyX and AxyY are more abundant in the resistant isolate (spectral counts of 62 *vs* 21 for AxyX and 10 *vs* 2 for AxyY in the results presented in Figure 1). However, the difference is not as pronounced as found for Axc and more accurate quantitative proteomics experiments have to be performed to validate these data.

Mutations in TetR-like repressors have been linked to differences in carbapenem resistance 53. For example, a 162 bp deletion in *axyZ* has been identified in certain carbapenem resistant strains ^54^. However, even though AxyZ_Gly29 leads to higher expression of *axyY*, resulting in higher MIC values for aminoglycosides, fluoroquinolones and tetracyclines, no correlation between *axyY* expression and meropenem resistance was observed ^52^. In line with this observation, a deletion of *axyY* in several *A. xylosoxidans* strains is reported to result in only a modest increase in the susceptibility towards carbapenems ^47^. Taken together, it is likely that AxyXY-OprZ per se contributes little if any to the meropenem resistance phenotype of our clinical isolate. Instead, our data show a critical role for Axc and suggest that *axc* expression is regulated by AxyZ. If this is indeed the case, we postulate that the increase in the meropenem MIC for the ACH-CF-911_V29G_ strain ^52^ is accompanied by increased Axc levels.

From the clinical perspective, the development of resistance to meropenem within days following meropenem treatment is remarkable. Previous longitudinal analyses of different *A. xylosoxidans* isolates from one patient have revealed large phenotypic and genetic differences, for example in the resistance towards different classes of antibiotics but they were generally performed over longer time periods ^35,53,54^. Such changes are believed to be the result of adaptive evolution of the initial strain which infected the patient, but there is also evidence that genetically different strains of *A. xylosoxidans* can co-exist within the same chronically infected individual ^55^. Though we cannot exclude a co-infection, it is likely that the mutation in *axyZ* in our case occurred during treatment.

Finally, from a diagnostic point of view, the presence of *Axc* in both the sensitive and resistant *A. xylosoxidans* isolates complicates straightforward detection by molecular methods in the future and warrants detection based on protein abundance levels. Moreover, in addition to the now well-established application of MALDI-ToF-MS for bacterial species identification, more elaborate mass spectrometry-based platforms clearly have potential for the detection of resistance and virulence proteins ^23,56^, as exemplified by the identification of Axc in meropenem resistant *A. xylosoxidans*.

## Methods

All procedures performed in studies involving human participants were in accordance with the ethical standards of LUMC Medical Ethical Committee and with the 1964 Helsinki Declaration and its later amendments or comparable ethical standards.

### Materials

MilliQ water was obtained from a Q-Gard 2 system (Merck Millipore, Amsterdam the Netherlands). Acetonitrile of LC-MS grade was obtained from Biosolve (Valkenswaard, the Netherlands). Porcine trypsin was purchased from Promega (Madison, WI). If not indicated otherwise, chemicals were from Sigma Aldrich (St Louis, MN, USA).

### Susceptibility profiling of clinical isolates

The minimum inhibitory concentrations (MICs) for different antibiotic compounds on the clinical isolates (Clinical IDs: M 14073954-7 (first isolate), M 14076260-2 (second isolate)) were initially determined using a Vitek-2 system (bioMérieux, Marcy-l’Étoile, France). Several colonies of plate grown cultures were inoculated and suspended in 0.45 % sterile physiological saline solution. Suspensions for testing had densities between of approx. 0.5 McFarland standards. The testing procedure was performed according to the manufacturer’s instructions.

Pure cultures of the susceptible and resistant isolates (AchroS and AchroR, respectively (Table 2)) were subjected to further susceptibility testing and used for the proteomic and genomic analyses. Etests (bioMérieux) were performed according to the recommendations from EUCAST (http://www.eucast.org/clinical_breakpoints/). EUCAST does not provide recommendations for the interpretation of MIC values or clinical breakpoints for *Achromobacter xylosoxidans*. Therefore, scoring was performed using locally developed protocols that are based on clinical breakpoints for other non-fermentative Gram-negative rods.

### Mass spectrometry-based proteomics

Cells were grown in BHI (Oxoid, Basingstoke, UK). Cells were collected by centrifugation (4000g, 5 min) from 1 mL cell culture and washed with phosphate-buffered saline (PBS, pH 7.4). Pellets were stored at −80° C until further use.

Two different proteomics experiments were performed. For the first, cell extracts from *A. xylosoxidans* strains (AchroS and AchroR, Table 2) were prepared in LDS (Lithium dodecyl sulphate) sample buffer (Novex, Thermo Scientific) and put at 95 °C for 5 minutes for cell lysis and protein extraction. Proteins were separated on Novex precast 4–12% Bis-Tris gels (Thermo Scientific, Waltham, MA, USA) with MOPS (3-(N-morpholino)propanesulfonic acid) running buffer (Thermo Scientific). After overnight staining using a Colloidal Blue Staining Kit (Thermo Scientific), destained gel lanes were processed into 31 slices per lane. Gel pieces were sequentially washed with 25 mM ammonium bicarbonate and acetonitrile. Reduction and alkylation were performed with dithiothreitol (DTT, 10 mM, 30 minutes at 56 °C) and iodoacetamide (IAA, 55 mM, 20 min at room temperature) respectively. Following several washes with 25 mM ammonium bicarbonate and acetonitrile, bands were overnight digested with trypsin (12.5 ng/μl in 25 mM NH_4_HCO_3_). Digest solutions were lyophilized and reconstituted in 0.5% trifluoroacetic acid (TFA). Nano-LC separation was carried out using a Ultimate 3000 RSLCnano System equipped with an Acclaim PepMap RSLC column (C18, 75 μm x 15 cm with 2 μm particles, Thermo Scientific) preceded by a 2 cm Acclaim PepMap100 guard column (Thermo Scientific). Peptide elution was performed by applying a mixture of solvents A and B with solvent A being 0.1% formic acid (FA) in water and solvent B 0.1% FA in 80 % acetonitrile (ACN). Peptides were eluted from the column with a multi-step gradient from 5% to 55% solvent B in 55 minutes, at a constant flow rate of 300 nl min^-1^. MS analysis was performed employing a maXis Impact UHR-TOF-MS (Bruker Daltonics, Bremen, Germany) in data dependent MS/MS mode in the *m/z* 150-2200 range. Ten ions were selected at a time, based on relative abundance and subjected to collision-induced-dissociation with helium as collision gas. A 1 minute dynamic exclusion window was applied for precursor selection.

For the second proteomics approach, in-solution digests were prepared as described previously ^23^. In short, following cell disruption proteins were solubilized in 50% trifluoroethanol (TFE). Subsequent reduction with DTT and alkylation with IAA were performed prior to overnight tryptic digestion. Samples were lyophilized and reconstituted in 0.5% TFA for injection. The nano-LC system and solvents were the same as in the above experiment, but using an Acclaim PepMap RSLC column (C18, 75 μm x 50cm with 2 μm particles, Thermo Scientific) with a 2 cm Acclaim PepMap100 guard column (Thermo Scientific). Peptides were eluted from the column with a multi-step gradient from 5% to 55% of solvent B in 180 minutes, at a constant flow rate of 300 nl min^-1^. MS analysis was carried out as described above.

### Mass spectrometry data export and spectral count analysis

Conversion of Bruker Impact files into mzXML format using CompassXport version 3.0.9, led to an initial total of 1,207,329 MS/MS and 444,565 MS/MS spectra, for the gel and total lysate based comparisons, respectively (in-solution total lysate digests were analysed twice and the data were merged). These spectra were searched using a concatenated forward and decoy strategy. The forward database was constructed from the 6386 unreviewed sequences from Uniprot for the organism *A. xylosoxidans* NH44784-1996 (November 2015) together with the cRAP contaminant sequences, as downloaded in January 2015. An in-house developed program, Decoy version V1.0.1-2-gfddc, that preserves homology, amino-acid frequency and peptide length distribution, was used with default flags to construct the decoy search space, which was concatenated to the forward sequences. The search against the resulting database was performed using Comet version 2014.02 rev.2, with precursor mass tolerance equal to 50 ppm and a fragment bin width of 0.05 Da, considering only fully tryptic digests with at most 2 missed cleavages. All cysteines were assumed carbamidomethylated, while methionine oxidation and N-terminal acetylation were regarded as variable modifications. The confidence of these results was assessed by means of Xinteract from the Trans Proteomics Pipeline suite version 4.8.0, retaining peptides longer than 6 amino acids and running in semi-parametric mode. A second in-house developed program, Pepxmltool version 2.5.1, was used to construct a protein quantification table based on spectral counting of only non-degenerate peptide (peptides mapping to a single protein) with corresponding q-value of at most 1%. Plots of the resistant versus susceptible quantifications of all proteins revealed for both methods a predominantly linear relationship, suggesting the applicability of an in-house developed program, Qntdiff version 0.1.1, to assign p-values to deviations from linear behaviour and provide lists of the most significantly differential protein expression levels between the two isolates. Custom-made Gnuplot scripts were written and run on Gnuplot version 4.6 patchlevel 4 to visualize differential proteins.

### Bacterial culture and genomic DNA preparation

AchroS and AchroR (Table 2) were cultured on trypcase soy agar plates (BioMérieux, Marcy l’Etoile, France), inoculated into liquid medium brain-heart-infusion (BHI) broth (Oxoid, Basingstoke, UK) and grown overnight (∼16-hrs) at 37 °C. Cells were harvested, washed with phosphate-buffered saline (PBS, pH 7.4), and genomic DNA extraction was performed using a phenol-chloroform extraction as previously described ^24^.

### Whole genome sequencing and SNP calling

Paired-end multiplex libraries were created as previously described ^25^. Sequencing was performed on an Illumina Hiseq 2000 platform (Illumina, San Diego, Ca, USA), with a read-length of 100 basepairs. High-throughput *de novo* assembly of sequenced genome was performed as previously described ^26,27^. The assemblies are then automatically annotated using PROKKA ^28^ with genus-specific databases from RefSeq ^29^. To identify single nucleotide polymorphisms (SNPs), the Illumina sequence data of the meropenem-susceptible *A. xylosoxidans* isolate (AchroS) was mapped on the assembled genome of the meropenem-resistant isolate (AchroR) using SMALT software (http://smalt.sourceforge.net/), after which SNPs were determined as previously described ^25^.

### PCR and heterologous expression of Axc

To corroborate the proteomics results and confirm results from the whole genome sequence analysis, the *axc* ORF, the *axc*-*axcR* intergenic region and part of the *axyZ* ORF were Sanger sequenced at a commercial provider (Macrogen, Amsterdam, the Netherlands). PCR products (for primers see Table 3) were sequenced using the same primers as those used for generating the product from genomic DNA isolated from the *A. xylosoxidans* isolates. We have submitted the Axc sequence to Genbank, ID MF767301.

**Table 3:**
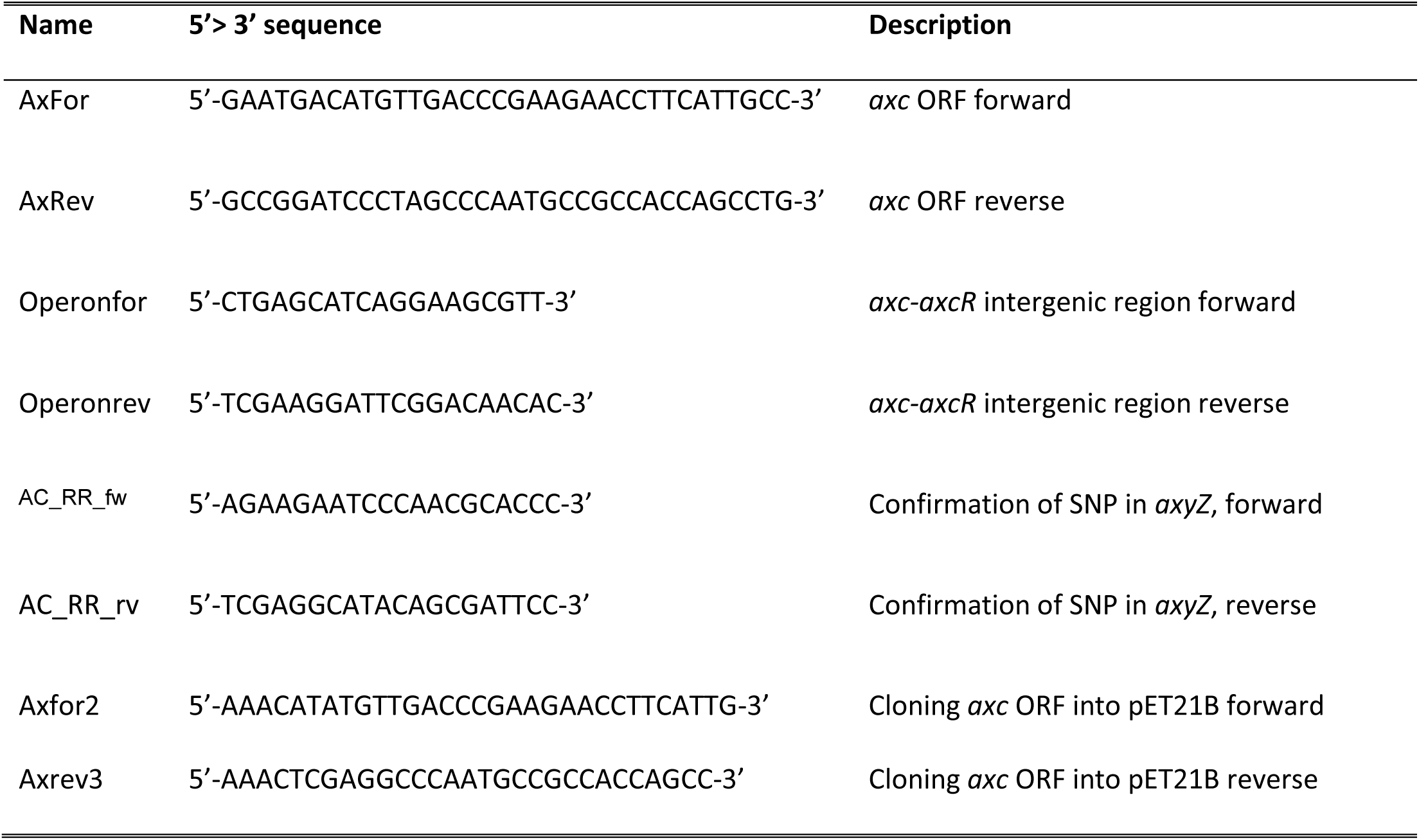
Primers used in this study. Axc: *<u>A</u>chromobacter <u>x</u>ylosoxidans* <u>c</u>arbapenemase. axcR: *a<u>xc</u>-* associated regulator. axyZ: TetR-type repressor of the *axyXY-oprZ* operon (36)

To construct an *E. coli* strain expressing *Axc* with a C-terminal 6xHis-tag from an IPTG (isopropyl β-D-1-thiogalactopyranoside) inducible promoter, the *axc* open reading frame was amplified using primers Axfor2 and Axrev3 (Table 3), using Accuzyme polymerase (GC Biotech, Alphen aan den Rijn, The Netherlands) and genomic DNA from *A. xylosoxidans* (AchroR) as a template. The amplified PCR product was digested using NdeI (Bioké, Leiden, The Netherlands) and XhoI (Roche, Almere, The Netherlands), and cloned into similarly digested pET-21b(+), yielding plasmid pET21B/Axc. The *axc* expression region was confirmed using Sanger sequencing. Expression of Axc-his6 was carried out in *E. coli* C43(DE3) ^30^ in Luria-Bertani broth (Affymetrix, Cleveland, OH, USA) with ampicillin (50 μg/mL) and 1mM IPTG (GC Biotech, Alphen aan de Rijn, the Netherlands) for 3 hrs at 37 °C and verified by immunoblotting using anti-His antibody (Agilent Technologies, Santa Clara, CA, USA).

### Susceptibility testing of *E. coli* expressing Axc

Minimal inhibitory concentration (MIC) values for the carbapenems imipenem (Etest, BioMérieux) and meropenem (microbroth dilution) were established for *E. coli* C43/pET21B-Axc (strain JC107, Table 2) in the presence and absence of 1mM IPTG. For imipenem, cells were grown overnight in LB broth at 37 °C in the presence of ampicillin (50 μg/mL). The overnight cultures were diluted 1:100 in LB broth with ampicillin and grown to mid logarithmic phase (OD_600nm_∼ 0.5). Two hundred μl of bacterial culture was spread on LB-ampicillin (50 μg/mL) plates (with or without 1 mM IPTG) and an Imipenem Etest (BioMérieux) was applied. MIC values were determined after 24h incubation at 37°C. The meropenem MIC values were established by microbroth dilution. Bacterial cultures in logarithmic phase (OD_600nm_∼ 0.5) were diluted into LB-ampicillin medium to an OD_600nm_ of 0.05 in the presence or absence of 1mM IPTG and subsequently seeded in a 96-well plate. A two-fold serial dilution of meropenem (starting at 12.5 μg/mL) was made by adding equal amounts of meropenem (25 μg/mL) to the first row, from which a two-fold dilution series was made in the rest of the plate. Samples were investigated for growth by measuring the OD_600nm_ after 24 hrs incubation at 37 °C, while shaking. The MIC was the lowest concentration of meropenem at which no growth was observed. As controls for our assays, a carbapenem resistant *Klebsiella pneumonia*e clinical isolate (KPC; JC113) and an unrelated expression construct (JC108; which expresses PPEP-1 ^31^ in an IPTG-dependent manner in the same *E. coli* C43 background, Table 2) were included.

### Colorimetric imipenem hydrolysis assay

Overnight bacterial cultures were diluted 100 fold in LB-ampicillin (50 μg/mL) medium and grown to exponential growth phase at 37 °C while shaking. At the time of induction, the OD_600nm_ was determined, the cultures were split in two and 1mM IPTG was added to one of the cultures, followed by a further incubation for 3 hrs at 37 °C while shaking. At T=3h, the OD_600nm_ was determined and cells were harvested by centrifugation (4000 g, 5 min) and stored at −20 °C overnight. Cells were resuspended in water to yield equal densities based on measured OD_600nm_ values. Then, 7.5 μL of bacterial suspension was mixed with 25 μL of imipenem/phenol red/ZnSO_4_ solution (3 mg/mL imipenem, 0.35% (wt/vol) phenol red, pH 7.8, 70 μM ZnSO_4_) and incubated at 37 °C for 1 hr. Conversion of imipenem leads to a pH drop that can be visualized by the color change of the buffer from red to yellow ^32^. To quantify this effect, the UV-Vis spectrum was determined with a Nanodrop ND1000 spectrophotometer (Thermo Scientific) and the ratio between the absorption peaks at 431 and 560 nm was taken as a measure of imipenem hydrolysis.

### NMR imaging of imipenem conversion

All proton nuclear magnetic resonance (^1^H-NMR) experiments were performed on a 600 MHz Bruker Avance II spectrometer (Bruker BioSpin, Karlsruhe, Germany) equipped with a 5-mm triple resonance inverse (TCI) cryogenic probe head with a Z-gradient system and automatic tuning and matching. All experiments were recorded at 310 K. Temperature calibration was done before each batch of measurements ^33^. The duration of the π/2 pulses was automatically calibrated for each individual sample using a homonuclear-gated nutation experiment on the locked and shimmed samples after automatic tuning and matching of the probe head. The samples were prepared by adding 70 μL imipenem aqueous solution (5 mg/mL) to 280 μL milliQ water. This solution was mixed with 350 μL 75 mM phosphate buffer (pH 7.4) in water/deuterium oxide (80/20) containing 4.6 mM sodium 3-[trimethylsilyl] d4-propionate. Twenty μL of bacterial cell suspension were added and the sample was mixed. Samples were manually transferred into 5-mm SampleJet NMR tubes. The cell suspension samples were kept at 6 °C on a SampleJet sample changer while queued for acquisition. For water suppression, presaturation of the water resonance with an effective field of γB1 = 25 Hz was applied during the relaxation delay. A 1D-version of the NOESY (Nuclear Overhauser effect spectroscopy) experiment was performed with a relaxation delay of 4 seconds. A NOESY mixing time of 10 ms was used during which the water resonance was irradiated with the presaturation RF field. After applying 4 dummy scans, a total of 98,304 data points covering a spectral width of 18,029 Hz were collected using 16 scans. The Free Induction Decay was zero-filled to 131,072 complex data points, and an exponential window function was applied with a line broadening factor of 0.3 Hz before Fourier transformation. The spectra were automatically phased and baseline corrected.

### Bioinformatic analysis

Comparison of Axc with other class A beta lactamases was performed by multiple alignment using the Geneious 9.0 (Biomatters Ltd, Auckland, New Zealand)) software algorithm for Global alignment with free end gaps, cost Matrix Blosum62. The tree was then built using Jukes-Cantor genetic distance model with the Neighbor Joining tree build method.

### Data availability

Illumina raw reads were deposited at the European Nucleotide Archive (ENA). Study ID: PRJEB19781. Sample IDs: ERS1575148 (AchroR) and ERS1575149 (AchroS).

## Acknowledgements

This research was financially supported by The Netherlands organisation of scientific research (NWO, ZonMW grant number 50-51700-98-142).

## Author contribution statement

F.F., H.C.v.L., S.T.N., E.J.K. and P.J.H. designed the research. F.F., J.C., C.W.K, I.D., N.K., A.V., H.C.v.L. and P.J.H. performed experiments. F.F., A.A.H., J.C., C.W.K, W.K.S., M.G., N.K., T.D.L.. A.V. and P.J.H. analyzed data. F.F., J.C., W.K.S., H.C.v.L., E.J.K. and P.J.H. wrote the paper

Competing interests: The authors declare that they have no competing interests

